# Demography, class-structure and kin selection in continuous-time models

**DOI:** 10.1101/456012

**Authors:** António M. M. Rodrigues

## Abstract

The Wright-Fisher infinite island model and the neighbour-modulated approach to kin selection have enabled major advances in the understanding of social evolution in a demographic context. Due to structural assumptions, however, some important evolutionary problems are difficult to solve within the Wright-Fisher discrete-time framework. Although these major constraints are relaxed in the Moran continuous-time framework, a formal treatment of the mathematics of kin selection in continuous-time class-structured populations is still lacking. Here, I employ the neighbour-modulated approach to formalise key features of the kin selection theory in continuous-time infinite-island models. Next, I derive a general form of Hamilton’s rule to enable an inclusive fitness interpretation of social behaviours. I consider class-structure at the group and individual level, and I focus on conditional and unconditional phenotypes. I illustrate how the general theory can be applied to solve a wide range of biological problems. Finally, I show how a simple extension of the framework allows for the study of problems pertaining to the transmission of parental quality. I show that while inheritance of parental quality may either promote or inhibit selection on conditional helping behaviours, unconditional behaviours are invariant with respect to the fidelity of inheritance.

## Introduction

Kin selection theory provides a general framework for studying the adaptive evolution of behaviours that affect not only the fitness of their bearers but also the fitness of their bearers’ social partners [1-4]. Kin selection is elegantly encapsulated in Hamilton’s rule, –*c* + *br* > 0, which provides the condition for the evolution of social traits [1,5]. A behaviour that inflicts a fitness cost *c* on the actor may nevertheless evolve if the fitness benefit *b* enjoyed by the recipient, depreciated by the relatedness *r* between actor and recipient, is sufficiently high. In simple scenarios, the canonical form of Hamilton’s rule can be readily applied. More often, however, multiple ecological, demographic and genetic variables will influence the leading quantities in Hamilton’s rule. Understanding social behaviour in such complex scenarios is a very active area of research and remains a major challenge for behavioural and evolutionary ecologists [4,6-9].

The inclusive-fitness and the neighbour-modulated approach to kin selection have been the two most widely used theoretical tools to study the evolution of social traits [1,6,10-13]. The inclusive fitness approach is actor-centric and focuses on how a trait expressed by a focal actor affects not only her own reproductive success but also the reproductive success of her social partners [12,13]. By contrast, the neighbour-modulated approach is recipient-centric, and it focuses on how a set of actors, composed by the social neighbours of the recipient, affect the reproductive success of the focal recipient [10,12,13]. While the formulation of these two approaches differs, they are mathematically equivalent, and therefore they yield the same predictions about the evolution of social traits.

Despite this equivalence, the neighbour-modulated approach has been particularly important for solving kin selection problems that unfold in a rich demographic context [6,8,11,12,14-26], where an inclusive-fitness approach is often harder to formulate. From the biological details of the problem, one can define the neighbour-modulated fitness of a focal recipient, and apply a well-studied mathematical optimisation method to derive the conditions under which social behaviours evolve [6,10,11]. In principle, we can then conceptualise these conditions in the form of Hamilton’s rule (e.g. [15-18]). In practice, however, not all studies obtain Hamilton’s rule, and when Hamilton’s rule is derived, it is often circumstantial and inconsistent between studies. Thus a general formulation of Hamilton’s rule that holds under very general ecological and demographic conditions is still lacking.

The most widely used framework to model demography has been the Wright-Fisher infinite island setting [27], in which populations are subdivided into several subpopulations connected by dispersal [15,19-26,28-32]. In part, the success of the Wright-Fisher infinite-island framework lies in being able to accommodate a considerable amount of ecological and demographic detail and still deliver analytical solutions. However, the Wright-Fisher framework also carries some structural assumptions that often limit the range of biological problems that can be addressed (see Rodrigues and Kokko [9] for a review). Habitat saturation is perhaps one of the most salient structural limitations of the Wright-Fisher framework, as most natural populations evolve in unsaturated habitats where empty sites are common.

A more recent breed of continuous-time models eases some of the most restrictive assumptions of the Wright-Fisher framework. More specifically, continuous-time models allow for the analysis of kin selection in unsaturated environments and account for ecological feedbacks, while keeping the analytic nature of the Wright-Fisher framework (e.g. Alizon and Taylor [33]; Wild et *al.* [34]; Rodrigues [35]). Alizon and Taylor, and Wild et *al.*, however, have relied on the more heuristic inclusive-fitness approach to kin selection. In the limited context of age-dependent social behaviour, I was able to derive Hamilton’s rule using both the inclusive-fitness and the neighbour-modulated approach, but I did not provide a formal treatment of the relationship between stochastic processes, which underlie the continuous-time models, and kin selection [35]. Therefore, a general formulation of the neighbour-modulated approach to kin selection in stochastic continuous-time models as well as a formal link between kin selection and the fundamental theory of stochastic processes remains unclear.

Here, I develop a general framework linking together the formal theory of kin selection and the fundamental theory of stochastic continuous-time models. First, I partition fitness into different components according to the properties of the demographic structure. Second, I establish a link between the fundamental theory of stochastic processes, kin selection and demography. Finally, I use the neighbour-modulated approach to provide a general account of kin selection in continuous-time models. Moreover, I consider between patch heterogeneity, where patches differ in quality (e.g. [15,26]), but also within patch heterogeneity, where individuals differ in quality (e.g. [17,31]). I consider behaviours that are expressed conditionally on the quality of both actor and recipient, but also behaviours that are expressed conditionally on the quality of patches, and behaviours that are expressed unconditionally. For each case, I show how to derive Hamilton’s rule using the neighbour-modulated approach, and how to obtain an inclusive fitness interpretation of the behaviour. I then illustrate how to apply the general framework to solve different biological problems, including the evolution of cooperation, virulence, and age-dependent helping. Finally, I show how a simple extension of this framework allows for the study of problems in which parental and offspring quality are correlated.

## Kin selection: General principles

I start off by providing a general formulation of fitness in class-structured populations [6,10,11,17,31,36]. In class-structured populations, individuals belong to different classes, in which class membership is associated with an individual’s contribution to the evolutionary process. Sex and age classes, for instance, are two of the most frequently observed class-structures in natural populations. However, class-structures pertaining to other forms of genetic or phenotypic differences among individuals are also common, such as class-structure that emerges from differences in the nutritional, physiological, or environmental state of individuals (e.g. [15,26,31]). In such general context, the neighbour-modulated fitness of a random individual in the population is given by

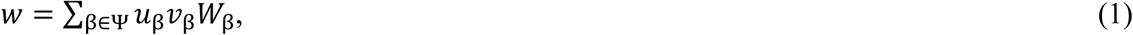

where: Ψ is the set of all possible classes; *u*_β_ is the frequency of individuals in the class *β*; *v*_β_ is the reproductive value of an individual in the class *β*; and *W*_β_ is the neighbour-modulated fitness of a random individual in the class *β* [6,10,11,31]. The fitness *W*_β_ of a random class-*β* individual is given by the individual’s genetic contribution to each class, denoted by *w*_β→η_, weighted by the reproductive value of each recipient class relative to the reproductive value of a class-*β* individual [6,10,31]. Thus, the fitness of a focal class-*β* individual is given by

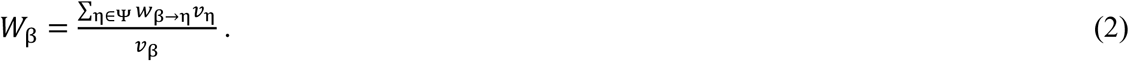

The set of class-specific reproductive successes, *w*_β→η_, form the elements of a square fitness matrix, where each row is associated with a recipient class, while each column is associated with a contributing class [6,10,17,31]. The fitness matrix is then given by

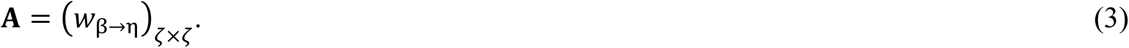

where *ζ* is the total number of classes in the population. The frequency and reproductive value of individuals in each class can be directly calculated from the fitness matrix [6,10,17,31,36]. The frequency of individuals in each class is given by the right-eigenvector of matrix **A,** while the reproductive value of each individual is given by the left-eigenvector, with both eigenvectors corresponding to the leading eigenvalue [10,17,31,36].

The frequency and reproductive value of individuals in each class can also be formulated as a dynamical system (e.g. [37]). Let **u** = (*u*1,*u*2,…,*u*ζ)ζ 1 be the column vector whose elements give the frequency of individuals in each class at any given time. The change over time in the frequency of individuals in each class is described by a dynamical system given by

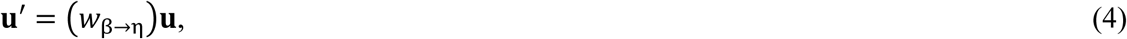

where **u**′ is the frequency of individuals in each class in the next time period. The changes in reproductive value over time obey to a similar system of equations, which is given by

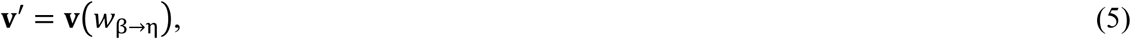

where **v** = (*v*_1_,*v*_2_,…,*v*_ζ_)_1 ζ_ is the row vector whose elements give the reproductive value of each individual in each class. Like the right- and left-eigenvector results, the solution of these systems of equations gives the asymptotic frequency and the reproductive value of individuals in each class (e.g. [37]).

## Kin selection and demography

My aim is to provide a general formulation of the fitness effects of a social trait *z* in a demographic context assuming vanishingly small genetic variation in the population [6,10,36]. I consider an infinite island model so that dispersal between patches is random and uniformly distributed [27,28]. I consider class-structure at the patch level, in which patches may be in different demographic states, but also class-structure at the individual level, in which individuals within each patch may be in different states. I build upon recent work that has sought to understand the evolution of social traits when patches vary in quality (e.g. [15,17,31]), as well as when individuals within each patch vary in quality (e.g. [17,31]).

### Fitness

To derive a fitness function in subdivided populations, I follow the formulation given in Rodrigues and Gardner [17], which considers both variation in the demographic state of patches across the population (i.e. between-patch heterogeneity) but also variation in individual quality within each patch (i.e. within-patch heterogeneity). I partition fitness into different additive components along three main dimensions. First, I consider reproductive success according to whether this is achieved through a philopatric or dispersed component, which I denote by the superscripts ‘ϕ’ and ‘δ’, respectively. Second, I consider class-structure at the patch level, in which patches are classified according to their state σ ∊ Ω, where the set Ω comprises all the possible patch demographic states, and where the state of patches can be defined according to different environmental or demographic variables, such as resource-availability [15,17], patch size [26], or age composition [35]. Finally, I consider class-structure within each patch, in which each individual is classified according to its quality *ρ* ∊ Ω_σ_, where the set Ω_σ_ comprises all the possible individual qualities present in type-*σ* patches, and where an individual’s quality can be defined, for instance, according to its future fecundity [31], maternal rank [17], or age [35]. I can then write the fitness of a focal class-*ρ* individual in a type-*σ* patch as

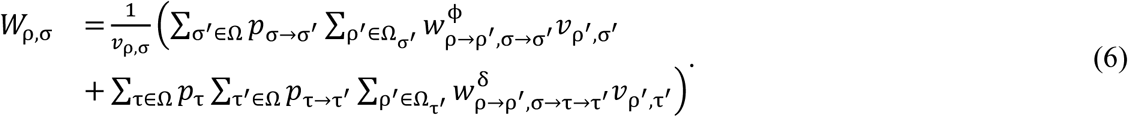

where: *p*_σ→σ′_ (or *p*_τ→τ′_) is the probability that a type-*σ* (or type-*τ*) patch becomes a type-*σ*′ (or type-*τ*′) patch; 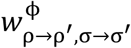 is the philopatric component of the reproductive success of a focal class-*ρ*′ individual when it produces class-*ρ*′ individuals associated with the demographic transition *σ*→*σ*′; *v*_ρ′,σ′_ (or *v*_ρ′,τ′_) is the reproductive value of a class-*ρ*′ individual in a type-*σ*′ (or type-*τ*′) patch; *p*_τ_ is the frequency of type-*τ* patches in the population; and 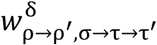is the reproductive success of a class-*ρ* individual in a type-*σ* patch when it produces class-*ρ*′ individuals that disperse and arrive at type-*τ* patches that become type-*τ*′ patches.

I assume that social interactions unfold within the local population, and therefore, I write down the fitness of a random individual in the local patch as

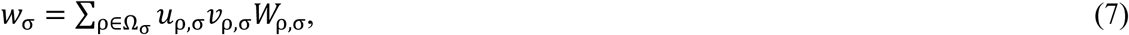

where the frequency of type-*ρ* individuals in a type-*σ* patch is given by *u*_ρ,σ_ = *p*_σ_*n*_ρ,σ_/∑_τ∊Ω_ *n*_τ_*p*_τ_, in which *n*_ρ,σ_ is the number of type-*ρ* individuals in a type-*σ* patch, and *n*_τ_ is the total number of individuals in a type-*τ* patch. The fitness of a random individual in the population is then given by

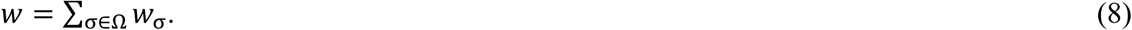

### Behaviour conditional on individual quality

Let us now consider the evolution of a social trait whose average level of expression is given by *z*. My aim is to obtain the fitness effects of a slight increase in trait value. I focus on a focal recipient whose reproductive success is mediated by her own phenotype *x*_ρ,σ,_ encoded by the breeding value *g*_ρ,σ,_ and by the phenotype of its neighbours *Y*_α,σ,_ encoded by the breeding value *G*_α,σ_ [6,17]. Actors are class-*α* individuals, while primary recipients (c.f. [31]) are class *ρ* ∊ Θ individuals, in which Θ is the set that comprises all the classes that contain the primary recipients of the behaviour, which may include the actor’s class. To determine the effect of a slight increase in breeding value on the fitness of the recipients in a type-*σ* patch, I take the derivative of fitness with respect to the breeding value. This is given by

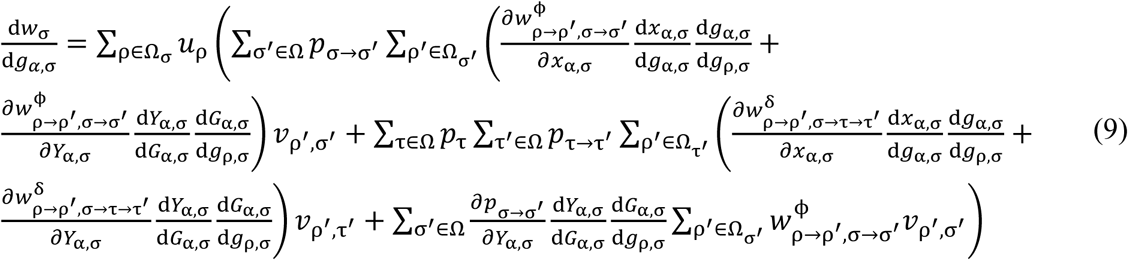

The partial derivatives are evaluated at the population’s average trait value *z*_ρ,σ,_ and they represent the effect of the phenotype on the reproductive success of each recipient (∂*w*/∂*x*), and on the demographic state of the patch (∂*p*/∂*x*). The derivatives of the actor’s breeding value with respect to the recipient’s breeding value, d*g*_α,σ_/d*g*_ρ,σ_ and d*g*_α,σ_/d*G*_ρ,σ,_ represent the coefficients of consanguinity between the actor and the recipients, denoted by *g*_αρ,σ,_ and the coefficient of consanguinity between the actor and herself, denoted by *g*_α•,σ_ [38]. The derivatives of the phenotypes with respect to the breeding values represent the mapping between the genotype and the phenotype, which we can set to one (i.e. d*x*/d*g* = d*Y*/d*G* = 1).

The coefficients of relatedness between the actor *α* and the recipient *ρ* in a type-*σ* patch, denoted by *R*_αρ,σ,_ is given by the ratio of the coefficient of consanguinity between the actor *α* and recipient *ρ* (*g*_αρ,σ_) and the coefficient of consanguinity between the actor *α* and herself (*g*_α•,σ_; [17,38]). Thus, *R*_αρ,σ_ = *g*_αρ,σ_/*g*_α•,σ_.

### Behaviour conditional on patch quality

So far, we considered cases in which individuals express a trait according to their quality. In some cases, however, information regarding individual quality may be unavailable, and consequently, individuals may have to resort to a strategy conditional on the quality of the local environment. Under this scenario, all individuals in the local patch are simultaneously actors and recipients, irrespective of their individual quality (e.g. [15,31]). As a result, the class of actors and recipients comprises all the classes present in the focal type-*σ* patch (Θ = Ω_σ_). The fitness effect of the trait is then given by

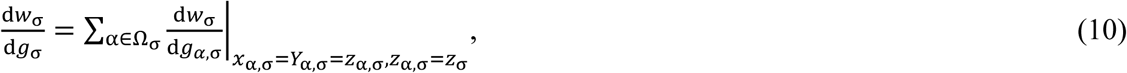

where all the partial derivatives are evaluated at the average level of the behaviour in type-*σ* patches (i.e. *x*_α,σ_ = *Y*_α,σ_ = *z*_α,σ_ and *z*_α,σ_ = *z*_σ_).

### Unconditional behaviour

In some cases, information regarding the state of the local patch may be unavailable to individuals, in which case social actors must express the behaviour unconditionally. In such cases, the behaviour must be considered across all patch types. The fitness effect of an unconditionally expressed behaviour is then given by

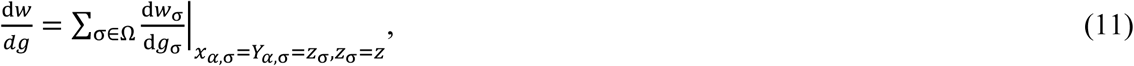

where the partial derivatives are all evaluated at the population’s average trait value (i.e. *x*_α,σ_ = *Y*_α,σ_ = *z*_σ_ and *z*_σ_ = *z*).

## Demography and continuous-time models

As shown in equation (9), kin selection depends on three key variables: first, the frequency of individuals in each class (i.e. *u*), which can be calculated from the frequency of each patch type (i.e. *p*; e.g. 33-35); second, the reproductive value of each individual (i.e. *v*); and finally, the relatedness among group mates (i.e. *R*). Here, I provide a formal link between the mathematics of stochastic continuous-time processes [39-42] and the mathematics of kin selection.

### Continuous-times models: general principles

Let us consider a discrete state space and continuous time, in which the stochastic process is defined by {*X*(*t*) ∊ Ω: *t* ∊ [0,∞)}, where *X*(*t*) is a discrete random variable with values defined by the set Ω, and where the argument *t* is continuous. The stochastic process {*X*(*t*) ∊ Ω: *t* ∊ [0,∞)} is denominated a continuous-time Markov chain (CTMC) and is characterised by the Markovian property, such that its current state depends only on the previous state and not on the states leading to the previous state. The random variable *X*(*t*) is described by the probability distribution {*p_σ_*(*t*)}*_σ_*_∊Ω_, where *p_σ_*(*t*) = Prob{*X*(*t*) = *σ*}. The transition probabilities between state *σ* and state *τ* are defined by *p_σ→τ_*(*t*, *t*_0_) = Prob{*X*(*t*) = *τ*|*X*(*t*_0_) = *σ*}, *t*_0_ < *t*, and I assume that the transition probabilities are homogenous such that

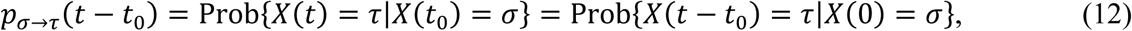

and therefore the transition probabilities depend only on the length of the time interval. The transition matrix is defined as **P**(*t*) = (*p*_σ→τ_(*t*))*_ξ_*_×*ξ*_, where *ξ* is the number of elements in the set Ω, and where the transition probabilities obey to the following property: ∑*_τ_*_∊Ω_ *p_σ_*_→*τ*_(*t*)= 1. The sequence of demographic states over time can be represented by the directed graph of the embedded Markov chain (see Figure 1 for an example). Let us define Dist(*σ*,*τ*) as the number of edges required to go from the demographic state *σ* to the demographic state *τ* given the shortest path between them. If we consider a Poisson process, for a sufficiently small time interval Δ*t* we have

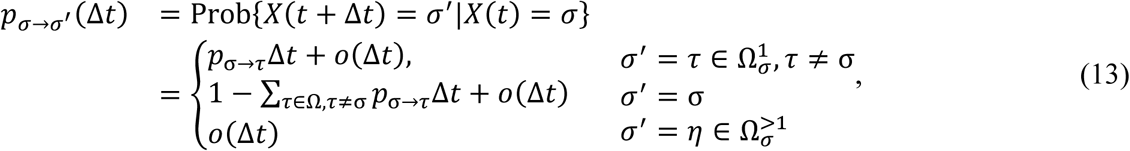

in which *o*(Δ*t*) – “little oh Δ*t*” – approaches zero more rapidly than Δ*t* when Δ*t* → 0, and where the set 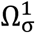 comprises the states that satisfy the condition Dist(σ,τ) = 1, whereas the set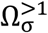 comprises the states that satisfy the condition Dist(*σ*,*η*) > 1.

**Figure 1.**
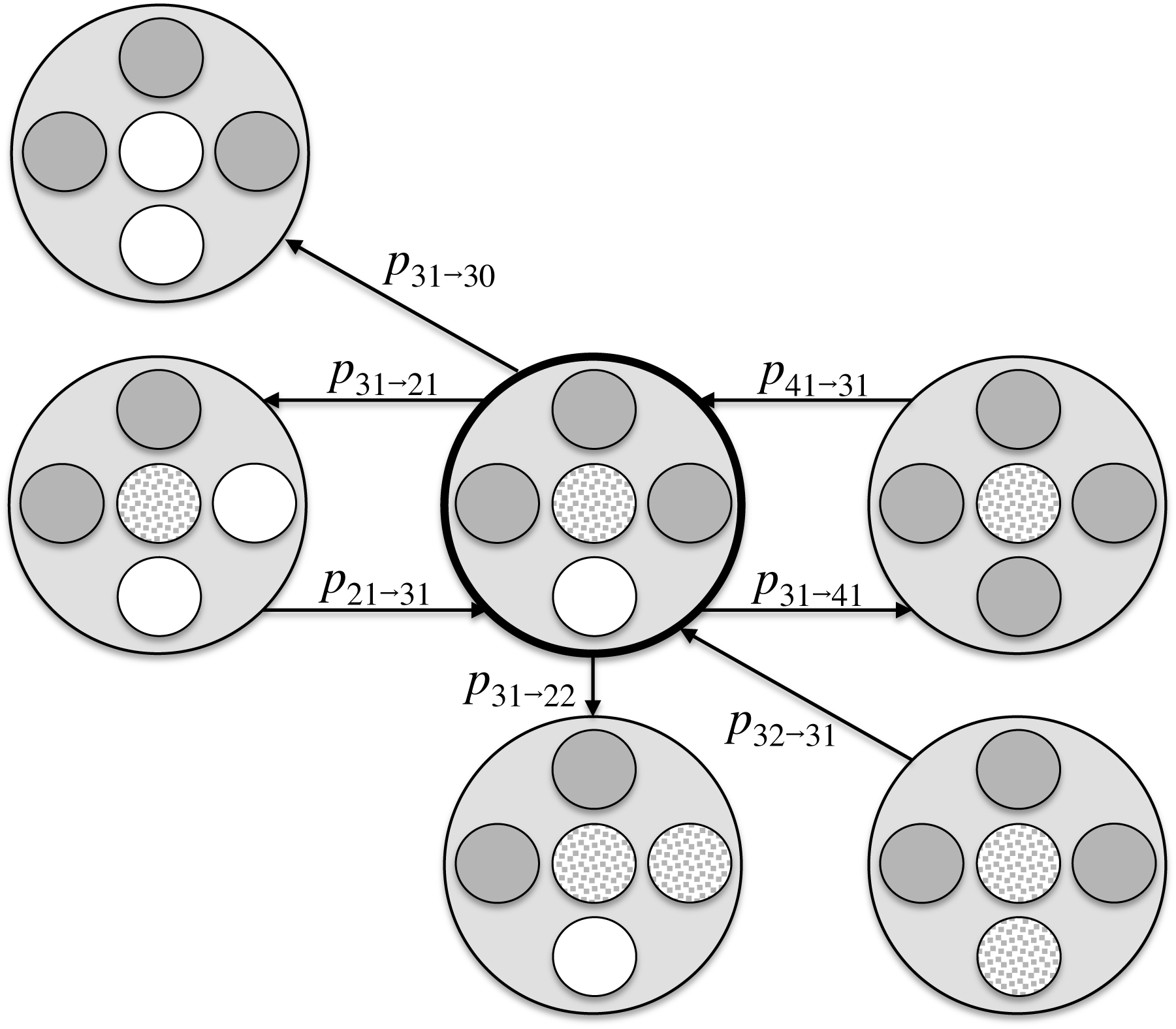
The directed graph of the embedded Markov chain. Larger circles represent the focal patch, while smaller circles represent breeding sites. Smaller grey circles represent sites occupied by class-1 individuals, while smaller spotted circles represent sites occupied by class-2 individuals. The indices represent the number of class-1 and class-2 individuals in the patch. Three different types of demographic transitions are represented with the corresponding demographic rates *p*. First, class-1 and class-2 individuals may give birth to class-1 individuals. Second, class-1 or class-2 individuals may die. Finally, Class-1 individuals may become class-2 individuals.

### Frequency of demographic states

As shown above, the fitness effects depend on the frequency of individuals in each class,which can be calculated from the frequency of patches in each state. Our CTMC is defined by equations (13), and therefore the probability that the Markov chain is in the state *σ* at time *t*+Δ*t* given by

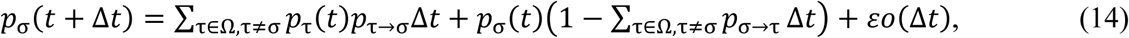

where εo(Δ*t*) comprises all the terms in o(Δ*t*) subtracting *p*_σ_(*t*) from both sides of the equation, dividing by Δ*t*, and taking the limit Δ*t* → 0, we get the forward Kolgomorov differential equations, which are given by

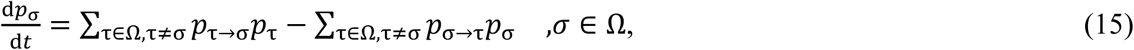

where the positive terms represent cases in which a patch in the demographic state τ ≠ σ becomes a type-*σ* patch, while the negative terms represent those cases in which a patch in the demographic state *σ* becomes a type-*τ* (≠ σ) patch, with *τ* ∊ Ω. Over time, the system of differential equations, for most cases, tends to an equilibrium state, which represents the asymptotic probabilities that the patch is in any of the given demographic states. We can represent this system of equations in a matrix form, by defining the generator matrix 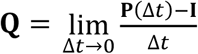, where **I** is the identity matrix. The forward Kolgomorov differential equations are then given by 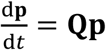 where **p**(*t*) = (*p*_σ_(*t*))_σ∊Ω_ is the vector of probabilities. The vector of probabilities provides a direct link to the infinite island model. The state of each island is described by the continuous-time Markov chain {*X*(*t*) ∊ Ω: *t* ∊ [0,∞)} defined above, and therefore, the probabilities **p** also give the frequency of each patch type in the population. The frequency of each patch type at equilibrium is given by the solution of the forward Kolgomorov differential equations. Note that the forward Kolmogorov differential equations in (15) recover the equations used in previous studies (i.e. Alizon and Taylor [33]; Wild et *al.* [34]; and Rodrigues [35]).

### Reproductive value

Second, the marginal fitness effects depend on the reproductive value of actors and recipients. Reproductive value measures the value of an individual according to her capacity to send copies of her genes to the gene pool of future generations [36,43,44]. Let us consider a very short time interval Δ*t* and how the reproductive value of class-*α* individuals in a type-*σ* patch changes over this time interval. From the fitness function (equation 6) and the transition probabilities of the CTMC (equations 13), the change in reproductive value is given by

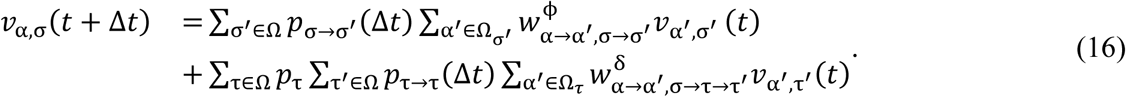

Expanding the right-hand side of this equation, and given that 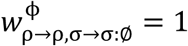, we obtain

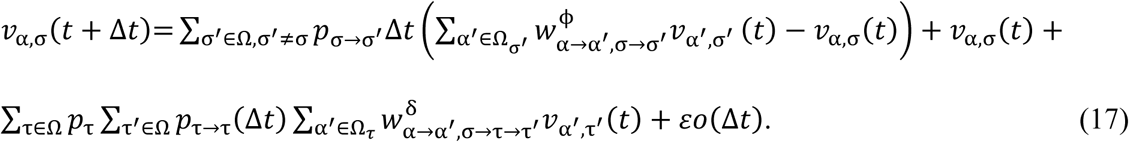

Subtracting *v*_α,σ_(*t*) from both sides of the equation, dividing by Δ*t*, and taking the limit Δ*t* → 0, we get the forward Kolgomorov differential equation for reproductive value, which is given by

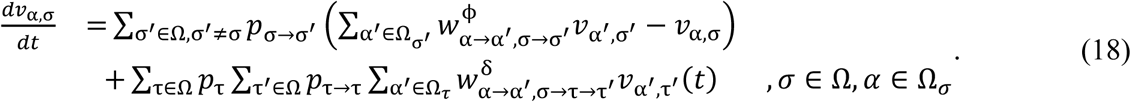

The system of differential equations describes the change in the reproductive value of each individual over time. At equilibrium, reproductive value will remain constant. We can then set the derivatives to zero, and solve the system of equations to obtain the reproductive value of each individual in a normal population (i.e. the mutant allele is neutral; [10,44]).

### Relatedness

Finally, the marginal fitness effects depend on the coefficient of relatedness among social partners, which gives the evolutionary measure of value an actor employs to evaluate social partners according to her genetic interests [1,2]. We can derive the coefficients of relatedness from the coefficients of consanguinity between social partners [16,17,38], and from the backwards probabilities associated with the demographic transitions in patch state (e.g. [15,17]). Specifically, the probability that given the demographic state *σ* the patch was in the demographic state *l* some Δ*t* time units ago is given by

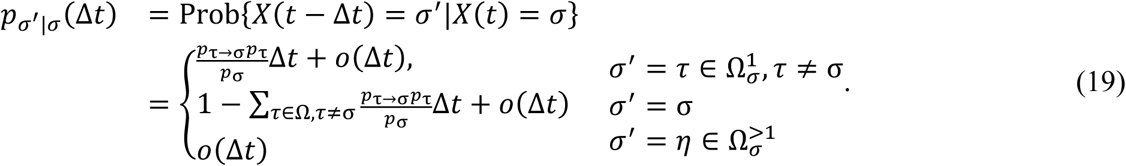

Thus, the change in the coefficient of consanguinity between class-*ρ* and class-*η* individuals in a type-*σ* patch over a short time interval Δ*t* is given by

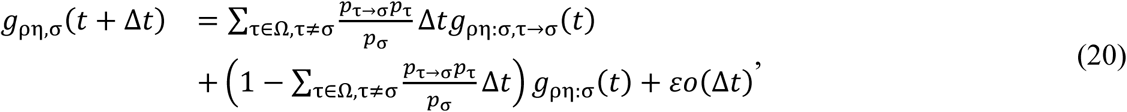

where *g*_ρη:σ,σ′→σ_(*t*) is the coefficient of consanguinity in a type-*σ* patch after the patch has transitioned from state *τ* to state *σ*. Note that if a type-*σ* patch has kept its demographic state over the Δ*t* time period, then the coefficient of consanguinity is simply *g*_ρη:σ_. Subtracting *g*_ρη:σ_(*t*) from both sides of the equation, dividing by Δ*t*, and taking the limit Δ*t* → 0, we get the differential equations for the coefficient of consanguinities, which is given by

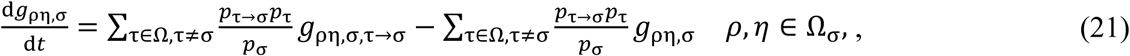

The system of equations can be solved to find the coefficients of consanguinity, which at equilibrium remains constant over time.

## Hamilton’s rule and inclusive fitness

Here, I derive a general formulation of Hamilton’s rule for a range of social traits. Hamilton’s rule enables an inclusive fitness interpretation of the behaviour, and therefore it establishes a formal link between the neighbour-modulated approach and Hamilton’s inclusive fitness theory [1,6,18]. I first analyse a helping behaviour that influences the fecundity of the actor and recipients, followed by the analysis of a helping behaviour that influences the survival of the actor and recipients. In Appendix A, I show how to extent the framework to analyse the evolution of a dispersal trait. For each trait, I consider behaviours that are expressed conditionally on the quality of individuals, but also traits that are expressed conditionally on patch quality as well as traits that are expressed unconditionally.

### General life-cycle

I consider a general model in which the fecundity rate of class-*ρ* mothers in type-*σ* patches is given by *f*_ρ,σ_. Offspring remain in the local patch with probability 1 – *d*_ρ,σ,_ and disperse to a random patch in the population with probability *d*_ρ,σ_. Offspring occupy empty breeding sites in type-*σ* patches at a rate *o*_σ_, and offspring who fail to acquire a breeding site die. Resident mothers die at a rate *m*_ρ,σ_. In addition, I consider that resident mothers may undergo changes in their reproductive, physiological, or social state. Such cases are represented by demographic rates of the form *a*_ρ→η_, which specify the rate at which class-*ρ* mothers become class-*η* mothers.

#### Demographic rates

Given these general model assumptions, let us consider key demographic rates. First, the demographic state of a focal type-*σ* patch may change whenever a mother dies. This is given by

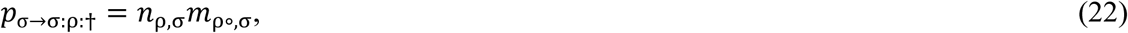

where: *m*_ρ∘,σ_ is the average mortality rate of class-*ρ* mothers in type-*σ* patches; and σ:ρ:† is the demographic state of a type-*σ* patch after a class-*ρ* mother has died. Second, the demographic state of a focal patch may change whenever an offspring occupies an empty breeding site. This is given by

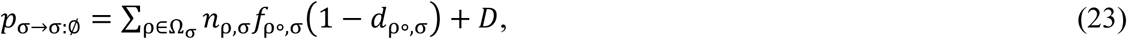

where: *f*_ρ∘,σ_ is the average fecundity of class-*ρ* individuals in the focal type-*σ* patch; *d*_ρ∘,σ_ is the average level of dispersal of class-*ρ* individuals in the focal type-*σ* patch; 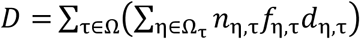 is the total number of immigrants that arrive at the focal patch, in which *d*_η,σ_ is the population’s average dispersal rate of class-*η* individuals in type-*τ* patches; and σ:Ø is the demographic state of a type-*σ* patch after a new offspring has occupied a vacant breeding site.

#### Reproductive success

We now need to determine the reproductive success of a focal mother associated with each of the demographic rates. First, let us consider the reproductive success of a resident mother when a class-*ρ* mother dies. If the mother is not a class-*ρ* mother, then her reproductive success is necessarily one. Otherwise, the reproductive success of a focal class-*ρ* mother is given by

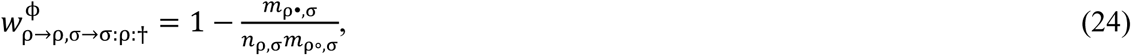

where 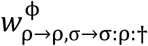 is the reproductive success of the class-*ρ* mother when a class-*ρ* mother dies in the focal type-*σ* patch, and *m*_ρ•,σ_ is the mortality rate of the focal class-*ρ* mother in the focal type-*σ* patch. Let us now consider the reproductive success of a resident mother when an offspring occupies a vacant breeding site. This has two additive components. First, the focal mother generates reproductive success via her own survival, which is necessarily one, i.e. 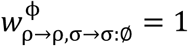. Second, the focal mother generates reproductive success if the offspring that occupies the empty breeding site in the local patch is her own. This is given by

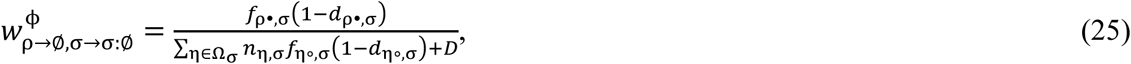

where: *f*_ρ•,σ_ is the fecundity of the focal class-*ρ* recipient in the focal type-*σ* patch; *d*_ρ•,σ_ is the probability of dispersal of the focal mother’s offspring; and 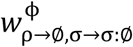 is the reproductive success of a focal mother via the production of philopatric offspring. Third, the focal mother generates reproductive success if her offspring occupy an empty breeding site in a foreign patch. This is given by

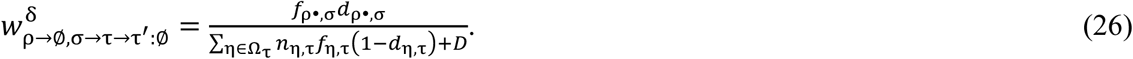

### Behaviour conditional on individual quality

#### Helping: fecundity effects

Let us consider a general helping trait that reduces the fecundity rate of the actor but raises the fecundity of the actor’s social partners, and that is conditionally expressed on the quality of the actor and recipients. I assume that actors are members of the class-*α*, and recipients are members of the class *β* ∊ Θ, where Θ is the set of the classes that include the primary recipients of the behaviour (i.e. those that receive the immediate benefits of the behaviour). The fecundity of a focal class-*ρ* recipient is given by *f*_ρ•,σ_ = *f*_ρ,σ_(*x*_ρ,σ,_*Y*_α,σ_), while the average fecundity of the class-*ρ* recipients is given by *f*_ρ∘,σ_ = *f*_ρ,σ_(*Y*_ρ,σ_,*Y*_α,σ_). All the other traits are set to the population’s average values. Because the behaviour influences the fecundity of each recipient, I focus on the demographic transitions and class-specific reproductive successes that depend on fecundity only, as outlined above. Replacing the expressions (22-26) in equation (9), we get

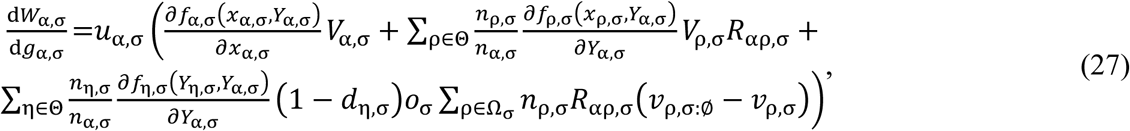

where *V*_α,σ_ (or *V*_ρ,σ_) is the reproductive value of an actor’s (or recipient’s) offspring, and *v*_ρ,σ:Ø_ is the reproductive value of a type-*ρ* individual when the focal type-*σ* patch accommodates a new breeder. The coefficient of relatedness among members of the same class *R*_αα,σ_ is the ‘whole-group’ coefficient of relatedness, which is given by *R*_αα,σ_ = 1/*n*_α,σ_+((*n*_α,σ_–1)/*n*_α,σ_)*r*_αα,σ,_ where *r*_αα,σ_ is the ‘others-only’ coefficient of relatedness [45]. The reproductive value of an offspring is given by

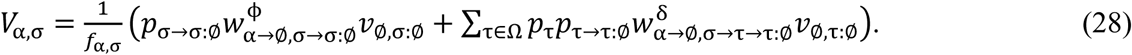

where *V*_Ø,σ:Ø_ (or *V*_Ø,τ:Ø_) is the reproductive value of an offspring that has taken up a breeding spot in a type-*σ* (or type-*τ*) patch. Note that ∂*f*_η,σ_(*x*_η,σ_,*Y*_α,σ_)/∂*Y*_α,σ_ is the effect of the class-*α* actors on the fecundity of a focal class-*η* recipient. Thus, the effect of a single class-*α* focal actor on class-*η* recipients is given by (*n*_η,σ_/*n*_α,σ_)(∂*f*_η,σ_/∂*Y*_α,σ_), which gives the inclusive fitness fecundity benefit *B_αη,σ_* provided to the class-*η* recipients by a focal class-*α* actor. Moreover, ∂*f*_α,σ_(*x*_α,σ_,*Y*_α,σ_)/∂*x*_α,σ_ is the effect of a class-*α* actor on her own fecundity, where the additive inverse gives the inclusive fitness fecundity cost *C*_α,σ_ to the actor. Replacing these variables in equation (27), we get Hamilton’s rule, which is given by

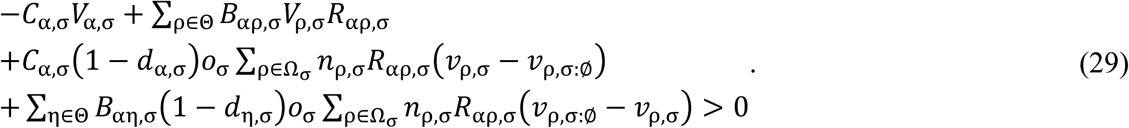

The left-hand side of this inequality immediately yields an inclusive fitness interpretation of the behaviour. A focal actor has *C*_α,σ_ fewer offspring, whose reproductive value is *V*_α,σ_. A class-*ρ* recipient enjoys a benefit that enables her to produce *B*_αρ,σ_ extra offspring, with each additional offspring leading to an increment *V*_ρ,σ_ in reproductive value, that must be depreciated by the relatedness *R*_αρ,σ_ between actor and recipient. Of the additional *B*_αη,σ_ offspring, a fraction 1 – *d*_η,σ_ remain in the local patch, which results in an empty site being occupied with probability *o*_σ_. The occupation of the vacant breeding site changes the reproductive value of all the *n*_ρ,σ_ class-*ρ* resident adults in the patch, who see their reproductive value change from *v*_ρ,σ_ to *v*_ρ,σ:Ø_. These changes in reproductive value must be depreciated by the relatedness *R*_αρ,σ_ between the actor and the recipients. Finally, the actor produces *C*_α,σ_ fewer offspring, who with probability 1 – *d*_α,σ_ would have stayed in the local patch, and with probability *o*_σ_ would have occupied a vacant breeding spot. This influences the demographic environment of the actor’s social partners, including herself, who see their reproductive value change from *v*_ρ,σ:Ø_ to *v*_ρ,σ_. These changes in reproductive value must be depreciated by the relatedness *R*_αρ,σ_ between the actor and recipients.

#### Helping: survival effects

Let us now consider a general helping trait that reduces the survival of the actor, denoted by *s*_α,σ,_ to improve the survival of the recipient(s), denoted by *s*_ρ,σ,_ in which *s*_α,σ_ = –*m*_α,σ_ and *s*_ρ,σ_ = –*m*_ρ,σ_. The survival of a focal class-*ρ* recipient is given by *s*_ρ•,σ_ = *s*_ρ,σ_(*x*_ρ,σ_,*Y*_α,σ_), whilst the average fecundity of class-*ρ* recipients is given by *s*_ρ∘,σ_ = *s*_ρ,σ_(*Y*_ρ,σ_,*Y*_α,σ_). All the other traits are set to the population’s average values. As above, we only need to consider the demographic rates and reproductive successes that directly depend on the survival of individuals. If we plug in the expressions (22-26) in equation (9), we get the following equation for the marginal fitness effects of the behaviour

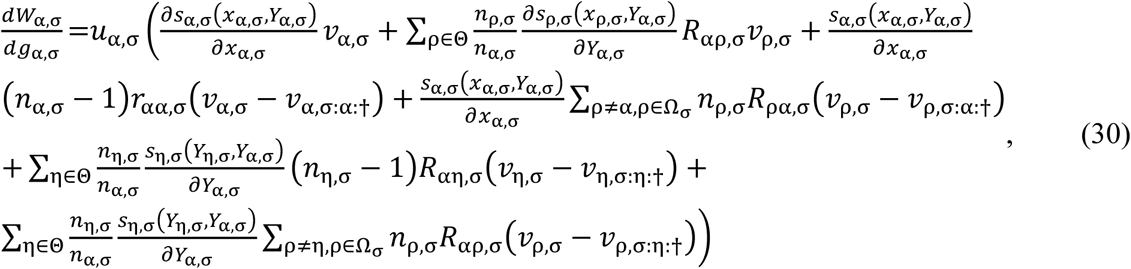

where *v*_α,σ_ is the reproductive value of the actor *α*, and *v*_α,σ:α:†_ is the reproductive value of a class-*α* individual after a class-*α* social partner has died. The same notation applies to the other variables.

Let us now consider the inclusive fitness survival costs and benefits. First, *s*_α,σ_(*x*_α,σ_,*Y*_α,σ_)/∂*x*_α,σ_ represents the effect of the class-*α* actor’s behaviour on her own survival, and consequently the additive inverse gives the inclusive fitness survival cost *C*_α,σ_ paid by the actor. Second, ∂*s*_ρ,σ_(*Y*_ρ,σ_,*Y*_α,σ_)/∂*Y*_α,σ_ represents the effects of the class-*α* actors’ behaviour on the survival of the focal class-*ρ* recipient, and consequently (*n*_ρ,σ_/*n*_α,σ_)(∂*s*_ρ,σ_/∂*Y*_α,σ_) gives the inclusive fitness fecundity benefit *B*_αρ_ provided by a class-*α* actor to class-*ρ* recipients. Replacing these variables in equation (30), we get Hamilton’s rule, which can be written as

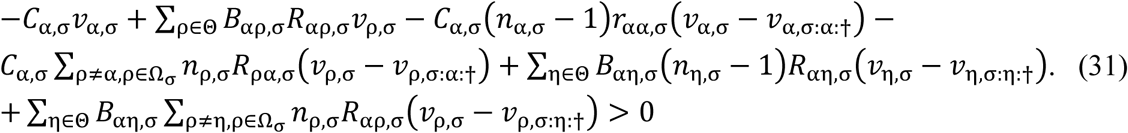

The left-hand side of this inequality immediately yields an inclusive fitness interpretation of the behaviour. An actor pays a survival cost *C*_α,σ_ and consequently loses all of her reproductive value *v*_α,σ_. A class-*ρ* recipient enjoys a benefit *B*_αρ,σ,_ which leads to an increment *v*_ρ,σ_ in her future reproductive value, an increment that must be depreciated by the relatedness *R*_αρ,σ_ between the actor and the recipient. The death of the actor has an impact on the other *n*_α,σ_ – 1 class-*α* individuals, who see their reproductive value change from *v*_α,σ_ to *v*_α,σ:α:†_. These changes in reproductive value must be depreciated by the relatedness *r*_αα,σ_ between the actor and her social and class partners. The death of the actor also impacts individuals in other classes, who see their reproductive value change from *v*_ρ,σ_ to *v*_ρ,σ:α:†_. Moreover, the additional survival of a class-*η* individual has an impact on the other *n*_η,σ_ – 1 class-*η* individuals, who see their reproductive value change from *v*_η,σ:η:†_ to *v*_η,σ_. These changes in reproductive value must be depreciated by the relatedness *R*_α*η*,σ_ between the actor and the class-*η* recipients. Furthermore, the additional survival of class-*η* individuals has an impact on individuals in other classes, who see their reproductive value change from *v*_ρ,σ:η:†_ to *v*_ρ,σ_. These changes in reproductive value must be depreciated by the relatedness *R*_αρ,σ_ between the class-*α* actor and the class-*ρ* recipients.

### Behaviour conditional on patch quality

The marginal fitness effects when behaviour is conditional on the state of the patch, but not on the state of individuals, follows immediately from equation (10). We now consider that all individuals are actors and recipients, irrespective of their class. Thus, the set of primary recipients of the behaviour is equal to the set of all classes present in the focal patch, i.e. Θ = Ω_σ_. In addition, the partial derivatives are now evaluated at the average trait value for the type-*σ* patch, which is given by *z*_σ_. Thus, *x*_α,σ_ = *Y*_α,σ_ = *z*_α,σ_ and *z*_α,σ_ = *z*_σ_, as shown in equation (10). The inclusive fitness interpretation is similar to the one given above for behaviours expressed conditionally on the quality of the individuals.

### Unconditional behaviour

Unconditional traits are averages of the marginal effects occurring at the different patch types. Hamilton’s rule follows immediately from equation (11). The marginal fitness effects must now be added together and the partial derivatives are evaluated at the population-wise average levels. Thus, *x*_σ_ = *Y*_σ_ = *z*_σ_. The inclusive fitness interpretation is similar to the one given above for behaviour expressed conditionally on the quality of the individuals.

## Applications of the general framework

Here, I analyse several examples to illustrate how the framework developed above can be employed in the study of kin selection problems. In particular, I focus on three studies that use the continuous-time framework but apply the inclusive-fitness method to kin selection to investigate the adaptive evolution of social traits (Alizon and Taylor [33]; Wild et *al.* [34]; and Rodrigues [35]). I show how the theory of Markovian processes is connected to these studies and how the general forms of Hamilton’s rule derived above can be readily applied to solve and analyse kin selection problems (see Appendix B for more details).

As we saw above from the general analysis of kin selection in a demographic context, fitness depends on three key variables: first, the frequency of patches in each of the demographic states; second, the reproductive value of each individual; and finally, the kin selection coefficients of relatedness between social partners. From the analysis of stochastic processes, we saw that these three quantities are calculated from the Kolmogorov differential equations defined by the system of equations (15), (18), and (21). These equations have precisely the same form of the equations used by Alizon and Taylor [33], Wild et al. [34], and Rodrigues [35] to calculate each of the key quantities. Thus, the Kolmogorov differential equations (15,18,21) provide a formal justification for the heuristics used in each of these three studies, and establish a link between their derivations.

Let us now consider Hamilton’s rule. Three key studies have employed the continuous-time framework to study the evolution of different social traits. Alizon and Taylor [33] studied a helping trait as a function of group size. Wild et *al.* [34] studied the evolution of parasite virulence. Finally, I studied the evolution of age-dependent social behaviour [35]. These studies generated a form of Hamilton’s rule to determine the selection acting on the traits of interest. In the appendix B, I show that each of these Hamilton’s rule are particular cases of the general formulations of Hamilton’s rule derived above. Thus, these general forms of Hamilton’s rule can be employed to analyse a wide range of evolutionary problems and unify different studies.

## Extensions of the framework: Transmission of individual quality

So far, in the general formulation of our problem, I assumed that there was no correlation between parental and offspring quality. In some cases, however, parents may transmit their quality to their offspring (e.g. [6]). Here, I show that these cases can be easily incorporated within the general framework. In particular, I consider a scenario where individuals are either of high- or low-quality and offspring can inherit parental quality to variable degrees of fidelity.

I assume that each patch has at most two resident breeders, who can be either high- or low-quality. High-quality (or low-quality) breeders give birth at a rate *f*_1_ (or *f*_2_) and die at a rate *μ*_1_ (or *μ*_2_). Offspring of high-quality (or low-quality) breeders become high-quality with probability *α* (or 1 – *β*), and low-quality with probability 1 – *α* (or *β*). Offspring remain in the natal patch with probability 1 – *d*_ij_, and disperse with probability *d*_ij_, where *i* is the quality of the mother and *j* if the quality of the mother’s social partner. When *j* = 0, then the quality-*i* mother is the sole resident breeder of the focal patch. Offspring settle in empty breeding spots at a rate *o*_l_ = *l*/*n*, where *l* is the number of empty breeding spots in the focal patch, and *n* is the total capacity of the patch (i.e. *l* = {0,1,2} and *n* = 2). I consider a helping trait that affects the survival of actor and recipient, and thefore Hamilton’s rule is given by inequality (31). In appendix C, I explain the methodology in detail and I use numerical methods to solve the model (e.g. [33-35]).

I find that in pure patches (i.e. patches occupied by breeders of the same quality), there is an increase in the potential for helping as the transmission of quality increases (Figure 2A). However, while in pure patches composed of low-quality individuals the potential for helping always increases with inheritance of quality, in pure patches composed of high-quality individuals the potential for helping decreases as inheritance of quality nears one (Figure 2A). This is because when transmission of quality increases, the density of high-quality immigrants also increases, which leads to a decrease in relatedness in patches composed of two high-quality individuals, but an increase in relatedness in patches composed of two low-quality individuals (Figure C1.A in Appendix C).

**Figure 2.**
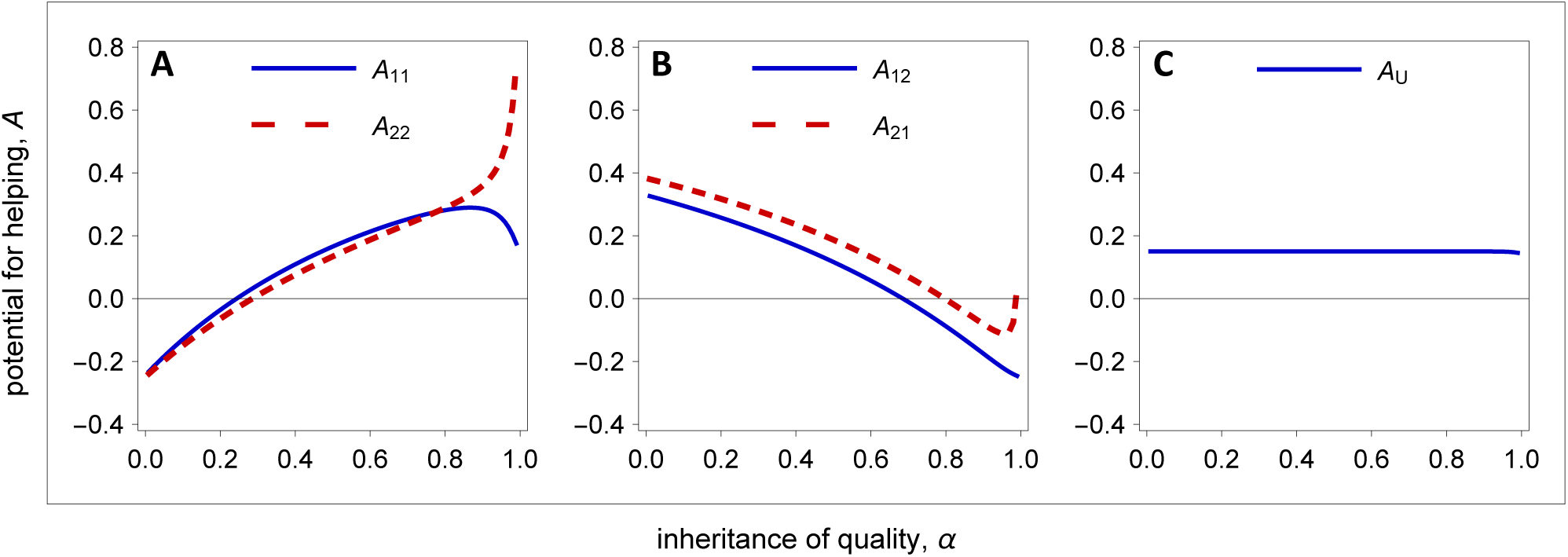
Potential for helping (*A*) as a function of the fidelity of quality inheritance (*α*). [A,B] Behaviour conditionally expressed on the quality of actor and recipient. [C] Unconditionally expressed behaviour. Parameter values: *β* = *α*, *f*_1_ = 2.2, *f*_2_ = 2.0, *μ*_1_ = *μ*_2_ = 0.3, *d*_10_ = *d*_20_ = 0, *d*_11_ = *d*_12_ = *d*_22_ = 1.

In mixed patches, there is a tendency for the potential for helping to decrease with the fidelity of inheritance (Figure 2B). This is because as the fidelity of transmission increases, the relatedness among social partners decreases (Figure C1.B). I also find that high-quality breeders are always selected to invest less in helping than low-quality breeders. This occurs because high-quality breeders have higher reproductive value than low-quality breeders (Figure C1.D). Finally, I find that as the fidelity of transmission approach one, the potential for helping of low-quality breeders increases with the fidelity of transmission (Figure 2B). This is because when the transmission of quality nears one, the reproductive value of low-quality breeders rapidly decreases (Figure C1.D).

We now turn the attention to the evolution of unconditional behaviour. I find that the potential for helping is independent of the fidelity of transmission (Figure 2C). While the potential for helping increases with the fidelity of transmission in pure patches, it decreases in mixed patches, such that average potential for helping depends little on the fidelity of transmission.

## Discussion

Here, I provided a framework connecting the formal theory of kin selection with the fundamental theory of stochastic continuous-time models. First, I derived a general expression of fitness in a demographic context. Second, I provided a link between the fundamental theory of stochastic processes, demography, and key kin selection variables. Finally, I derived Hamilton’s rule for major social traits. Specifically, I employed the neighbour-modulated approach to kin selection to provide a general account of inclusive fitness in continuous-time models. I partitioned the neighbour-modulated fitness of a focal recipient to account for population spatial structure, environmental and demographic variation across the population, as well as variation in individual quality within each patch. Starting with the neighbour-modulated fitness of a focal recipient, I then derived the marginal fitness effects for the expression of different kinds of behaviours. In particular, I considered behaviours that are expressed conditionally on an individual’s quality, but also behaviours that are expressed conditionally on patch quality as well as unconditionally expressed behaviours.

I showed how to calculate three key variables that mediate the fitness effects of a behaviour, namely: the frequency of patches in each demographic state; the reproductive value of each individual; and the kin selection coefficients of relatedness. I then obtained the expressions for the reproductive success of each recipient from the demographic transition rates that characterise continuous-time models. Given the fitness effect of a behaviour, the demographic rates, and the expressions of reproductive success, I derived a general formulation of Hamilton’s rule for a helping and a dispersal trait and the corresponding inclusive fitness interpretation. Next, I illustrated how this general framework can be used to study and unify a wide range of kin selection problems.

In a first instance, I derived a general expression for the fitness effect of a social behaviour. In the context of a Wright-Fisher infinite island model, Rousset and Ronce [14] derived a similar expression. However, their marginal fitness expression is narrower in scope and partitioned differently. Like my expression, they have partitioned marginal effects according to whether fitness is achieved through a philopatric or dispersed component, according to the demographic state of patches, and according to whether the marginal fitness effects emerge from changes in the reproductive success of individuals or in the demographic state of the focal patch.

However, their partition of the marginal fitness also differs from mine in significant ways. First, they have not considered variation in individual quality within patches as captured by my framework. Second, while they focused on changes in the demographic state of patches that emerge from alterations in patch size, my framework also consider other kinds of demographic changes that do not involve modifications in patch size. For instance, within my framework, changes in demography can occur through changes in the age composition of the group or in the infected state of group members. Finally, while their work focuses on the study of the marginal fitness effects, I extracted general formulations of Hamilton’s rule for major social behaviours from the marginal fitness effects.

I established a formal link between the neighbour-modulated approach and Hamilton’s inclusive fitness theory in a population with complex demographic and spatial structures. This link is significant for several reasons. First, the empirical testing of hypothesis and the design of experiments often involves an inclusive fitness argument and therefore the adoption of an actor-centric perspective, rather than then the recipient-centric perspective adopted in the neighbour-modulated approach [1,3,4,13,18]. Second, the inclusive fitness perspective provides a common ground that is able to connect not only theoretical and empirical thinking but also theoretical studies that employ different methodologies [18]. In particular, recent years have witnessed the proliferation of different methodologies to determine the intensity of selection operating on social traits, from inclusive fitness to multilevel selection. Hamilton’s rule and the inclusive fitness perspective provide a conceptual tool to unify and contrast results obtained from different methods.

Third, while in abstract terms theory suggests that inclusive fitness is as general as natural selection [13,46], in practice, a large number of studies have contested this idea (reviewed in [46]). My work shows that inclusive fitness holds under a wide range of conditions in the context of populations with complex demographic and spatial structures. Finally, there is the general conviction that the neighbour-modulated approach yields the same results than the inclusive fitness approach [12]. However, practitioners of the neighbour-modulated approach not always provide an inclusive fitness interpretation of the behaviour (e.g. 47,48]). When the inclusive fitness perspective is given, it is frequently inconsistent among different studies, with the interpretations of the behaviour varying significantly across studies (e.g. [49]). In some other cases, an interpretation made in one context breakdowns when other scenarios are considered (e.g. [15,21,22,26]). My study provides an unifying conceptual tool that integrates an increasingly disconnected body of work.

I derived a general form of Hamilton’s rule for a helping trait, under both fecundity and survival effects, and for a dispersal trait. From these general forms of Hamilton’s rule, I was able to immediately recover previous results that were obtained using the inclusive fitness method [i.e. 33-35]. In particular, I was able to recover results pertaining to the evolution of a helping trait under fecundity effects [33], the evolution of parasite virulence [34], and the evolution of age-dependent helping and dispersal [35]. The diversity of solutions implicit in the general forms of Hamilton’s rule suggests that it should be relatively easy to obtain results for many other social traits, but also that further explorations of my framework is likely to generate deeper insights concerning the fundamental forces driving the evolution of social traits.

I have established a formal link between stochastic processes and the equations that characterise three key quantities in kin selection models: patch frequencies, relatedness and reproductive value. In all three previous studies, Alizon and Taylor [33], and Wild et al. [34], and Rodrigues [35], the equations that characterise these three key quantities were obtained using heuristic arguments. Above, I was able to provide a formal link between the theory of stochastic processes and the equations for each of these quantities under very general demographic and ecological conditions.

The general formulation of the problem under consideration can be easily extended to consider cases in which parental quality is correlated with offspring quality. In such cases, the class of the offspring will depend on the quality of the parent, and this has an impact on the selection on helping behaviours. The fidelity of quality inheritance between parents and offspring has an impact in the demographic and genetic structure of the population, which may either promote or inhibit the evolution of cooperation. However, when I consider unconditional phenotypes, the fidelity of quality inheritance has little (if any) influence on the evolution of social behaviour. More generally, the impact of quality inheritance on the evolution of helping has received little attention, and therefore future studies should explore this problem further.

I focused on an asexually-reproducing and haploid species. However, extending the current framework to sexually reproducing populations is straightforward. This involves additional sex-classes in addition to other types of class-structures. In such cases, the calculation of reproductive value must take sex-structure into account by considering the genetic contribution of individuals to each sex class [16,17]. Moreover, the calculation of the coefficients of consanguinity involves the evaluation of the levels of inbreeding within each sex in addition to the coefficient of consanguinity between social partners [16,17].

I considered heterogeneity both between and within patches. In the examples provided above, heterogeneity between patches emerges because the size or composition of patches may vary owing to the life history of individuals. In some cases, however, differences in patch quality may be imposed owing to extrinsic forces, such as environmental changes [e.g. 15]. Such cases are easily handled within the framework developed in this study. In addition, I have considered behaviours expressed conditionally on the quality of the actor and on the quality of the recipient. However, we could consider other scenarios, such as when the behaviour is expressed conditionally solely on the quality of the recipients, in which cases one has to define the set of classes that enact the behaviours (e.g. Rodrigues and Gardner [31]).

## Competing interests

I have no competing interests.

## Data Availability

This study did not use any data.

## Competing Interests

I have no competing interests.

## Authors’ Contributions

AMMR designed and carried out the analysis of the model. AMMR wrote the manuscript. The author gave approval for publication.

## Funding

The author received no funding for this study.

## Research Ethics

This study did not require an ethical assessment.

## Animal Ethics

This study did not require an animal ethical assessment.

## Permission to Carry out Field Research

No permission to carry out field research was required.

## Acknowledgements

I thank Wolfson College, Cambridge for support and Behavioural Ecology Group, Cambridge for helpful discussions.

